# Genomic prediction of cereal crop architecture traits using models informed by gene regulatory circuitries from maize

**DOI:** 10.1101/2024.08.01.606170

**Authors:** Edoardo Bertolini, Mohith Manjunath, Weihao Ge, Matthew D. Murphy, Mirai Inaoka, Christina Fliege, Andrea L. Eveland, Alexander E. Lipka

## Abstract

Plant architecture is a major determinant of planting density, which enhances productivity potential for crops per unit area. Genomic prediction is well-positioned to expedite genetic gain of plant architecture traits since they are typically highly heritable. Additionally, the adaptation of genomic prediction models to query predictive abilities of markers tagging certain genomic regions could shed light on the genetic architecture of these traits. Here, we leveraged transcriptional networks from a prior study that contextually described developmental progression during tassel and leaf organogenesis in maize (*Z. mays*) to inform genomic prediction models for architecture traits. Since these developmental processes underlie tassel branching and leaf angle, two important agronomic architecture traits, we tested whether genes prioritized from these networks quantitatively contribute to the genetic architecture of these traits. We used genomic prediction models to evaluate the ability of markers in the vicinity of prioritized network genes to predict breeding values of tassel branching and leaf angle traits for two diversity panels in maize, and diversity panels from sorghum (*S. bicolor*) and rice (*O. sativa*). Predictive abilities of markers near these prioritized network genes were similar to those using whole-genome marker sets. Notably, markers near highly connected transcription factors from core network motifs in maize yielded predictive abilities that were significantly greater than expected by chance in not only maize but also closely related sorghum. We expect that these highly connected regulators are key drivers of architectural variation that are conserved across closely related cereal crop species.

**Article summary:** We used an approach typically used for breeding to infer the contributions of biological gene networks to plant architectural traits. We found that markers near genes belonging to smaller, specialized gene networks from maize could predict breeding values of leaf angle better than expected by chance for both maize and sorghum.

## Introduction

The increasingly pressing challenge of feeding the growing global population, estimated to reach nine billion by 2050, necessitates a significant increase in food production (Hunter et al. 2017). Against the backdrop of climate change, this challenge underscores a critical need to explore cutting-edge methods for crop improvement (Mohd Saad et al. 2022). Plant architecture has been an important target of selection in crop improvement and central to the huge gains in productivity from breeding seen throughout the past century. With inevitable decreases in arable farmland worldwide and shifting weather patterns, modern day crop improvement must adjust plant ideotypes for diverse environments and thus architecture traits remain a key target (Huang et al. 2022).

A core breeding objective to accommodate these challenges has been to increase planting density without compromising plant fitness. Consequently, modern high-density planting schemes for cereal crops have been successfully implemented, resulting in significantly increased productivity per unit area (Cao et al. 2022). In maize (*Z. mays*), this was realized through breeding efforts focused on architecture traits such as upright leaves, minimal branching, and tillering. Narrower leaf angle (LA) and reduced tassel branch number (TBN) not only have significant implications for crop management but also enable better light penetration into the lower canopy. This reduces competition for sunlight and hence optimizes photosynthetic efficiency. Such breeding efforts resulted in a higher number of plants per hectare and, consequently, a substantial increase in crop yields (Duvick 2005).

Plant architecture traits, including LA and inflorescence structure (e.g., TBN), exhibit high heritability across economically important crops (Sary et al. 2022; Upadyayula et al. 2006; Casa et al. 2008). This allows the opportunity to employ genomic prediction (GP) models that utilize signatures of genetic architecture captured by genome-wide marker sets to obtain accurate genomic estimated breeding values (GEBVs) for plants that have not been phenotyped (Bernardo 1994; Meuwissen, Hayes, and Goddard 2001) and therefore accelerate genetic gain. The most widely-used statistical models for GP (reviewed in de Los Campos et al. (2013) have been shown to outperform competing marker-assisted selection approaches that only consider the effects of single, large effect genes (Meuwissen, Hayes, and Goddard 2001; Bernardo and Yu 2007; Heffner et al. 2010). In addition to its well-established contribution to modern plant and animal breeding programs (reviewed in (Van Eenennaam et al. 2014; Crossa et al., n.d.), GP models that account for gene families or other genomic features identified in prior studies can be used to make inferences on their contribution to the overall genetic architecture of the studied trait. For example, Turner-Hissong et al. (2020) used a series of fine-tuned GP models to infer the contributions of several pathways to free amino acids composition in dry *Arabidopsis* seeds.

Prior studies investigated the use of certain genomic features to reduce genome-wide marker sets, e.g., accessible chromatin regions, capturing a significant portion of the phenotypic variation (Rodgers-Melnick et al. 2016; Parvathaneni et al. 2020). A recent study by Bertolini et al. (2024) demonstrated the power of fine-tuning genotype-to-phenotype models with biological information derived from transcriptional networks, which facilitated the identification of small effect genetic loci associated with LA and TBN in maize (Bertolini et al. 2024). This approach defined transcriptional circuitries in specific developmental contexts (i.e., tassel and leaf development) (**Figure 1**, Left Panel) by inferring 1) gene co-expression networks to identify groups of genes with similar expression patterns and 2) gene regulatory networks to predict transcription factor regulation of gene targets. This study revealed that gene sets derived from specific developmental networks could explain a significant portion of the narrow-sense heritability (*h^2^*) for LA and TBN, which suggests these gene sets form part of the genetic architecture for these traits (Bertolini et al. 2024). Therefore, we expect that these sets of network genes should similarly produce reasonably high predictive abilities in GP models.

**Figure 1:**
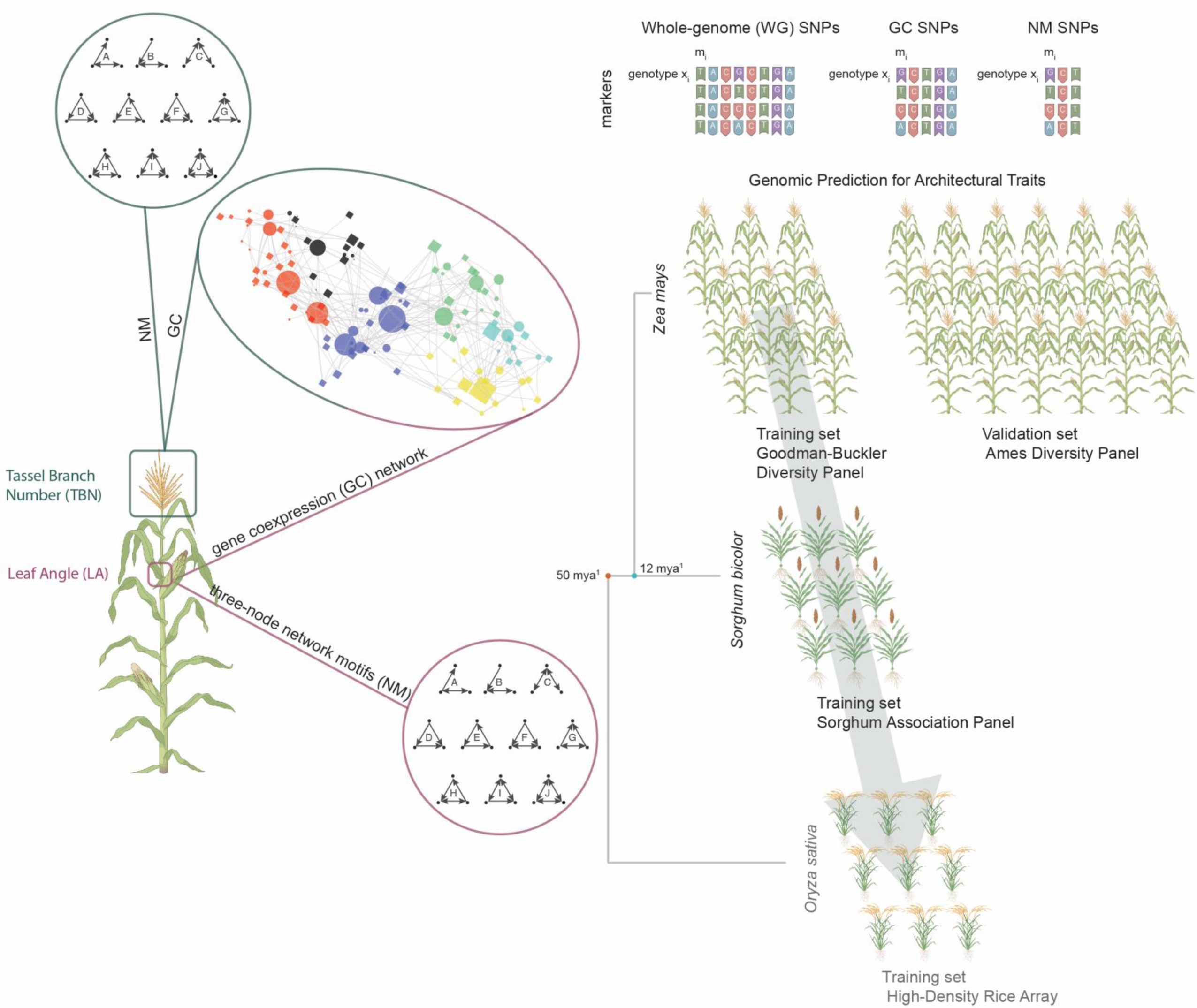
Experimental scheme of genomic prediction using transcriptional networks. The left panel represents the identification of transcriptional networks from tassel primordia and the ligular region (Bertolini et al. 2024). The ball-and-stick diagram, enclosed within the red-green ellipse, represents the six selected modules from the co-expression networks generated using expression data from both tissue types (Bertolini et al. 2024). The red and green sets represent the tissue-specific three-node network motifs with an edge number of three or more (from A to J). The right panel illustrates the genomic prediction approach used in this study. It includes three marker sets (WG, GC, NM), each comprising markers with minor allele frequency (MAF) less than 0.05 and then markers with MAF greater than 0.05. Genomic prediction models were trained in the maize Goodman-Buckler Diversity panel and validated in the Ames panel. The gray arrow indicates the cross-species translation using markers located near sorghum and rice orthologs. Blue and orange dots represent phylogenetic distance (million years ago) among the species (Swigonová et al. 2004).

The purpose of our study was to use GP to determine the contribution of biological gene networks to the genetic architectures of LA and TBN, two agronomically important architecture traits in maize. We also tested the ability to translate gene network-based information from maize to other cereals including sorghum (*S. bicolor*), which is closely related to maize, and rice (*O. sativa*), a more distantly related cereal. We considered four prediction scenarios (**Figure 1**, right panel): 1) where we predicted GEBVs within the maize diversity panel from (Flint-Garcia et al. 2005); 2) where the maize diversity panel in 1) was the training set and a larger maize diversity panel from (Romay et al. 2013) was the validation set; 3) where the predictive abilities for LA using sorghum orthologs of core network genes from maize were assessed in a sorghum diversity panel (Casa et al. 2008); and 4) where a procedure similar to 3) was conducted on a rice diversity panel (McCouch et al. 2016). For all scenarios, we compared the predictive abilities of markers proximal to network gene sets to those from a genome-wide marker set, as well as an empirical distribution of prediction accuracies from randomly selected subsets of markers.

## Materials and Methods

### Genotypic data

We analyzed two well-studied maize diversity panels: the Goodman-Buckler Diversity Panel (Flint-Garcia et al. 2005) and the Ames Panel (also known as the North Central Regional Plant Introduction Station Panel; Romay et al. 2013). We also analyzed the Sorghum Association Panel (SAP; Casa et al. 2008) and the High-Density Rice Array (HDRA; McCouch et al. 2016). Genotyping By Sequencing (GBS) data for the maize diversity panels were downloaded from Panzea (www.panzea.org) and processed following the methodology outlined in (Bertolini et al. 2024). Specifically, genomic coordinates were uplifted to the maize reference AGPv4 (Jiao et al. 2017), indels and non-biallelic markers were filtered out, and missing data were imputed using the nearest neighbor method (Money et al. 2015). Single-nucleotide polymorphisms (SNPs) with a minor allele frequency (MAF) less than 0.01 were also discarded. The SAP GBS data (Bouchet et al. 2017) were downloaded from the Dryad Digital Repository (doi:10.5061/dryad.gm073). The HDRA genotypes data were downloaded from Rice Diversity (www.ricediversity.org). All genotype data were then employed to subset markers based on specific gene network subsets and different MAF cut offs for further analysis.

### Gene module information

We used the gene-coexpression (GC) networks and three-node network motifs (NM) related to tassel branching and ligule development in maize from the Bertolini et al. (2024) study. Markers within genomic coordinates of these two gene sets were selected based on genomic windows defined as within 2 kb from the transcription start site (TSS) and the transcription termination site (TTS) (see Bertolini et al. (2024) for further details). The maize GC and NM genes were translated to sorghum and rice using syntenic-orthologous gene information retrieved from Zhang et al. (2017). Due to larger LD blocks relative to maize (Morris et al. 2013), the sorghum and rice genomic regions were extended by 10 kb from the TSS and TTS.

### Phenotypic data

We used phenotype data for LA and TBN published in (Bertolini et al. 2024) for our analysis. These data were from 231 lines of the Goodman-Buckler diversity panel and 1,064 lines of the Ames panel. As described in (Bertolini et al. 2024), these data were grown in a randomized complete block design (RCBD) between 2018-2021. Sorghum leaf angle phenotype data were previously collected for 296 individuals from the Sorghum Association Panel (SAP) (Casa et al. 2008), which were planted in a RCBD with two replications per location in 2010 and 2012. LA was measured from the leaf below the flag leaf, and two plants per replication were measured, using a protractor (Mantilla Perez et al. 2014). Similarly, LA phenotype data were collected from a rice diversity panel of 344 varieties (Huber et al. 2023) using a RCBD with four replicates. LA was collected at an early vegetative stage, between the second and third youngest leaves and the culm.

### GP model used

We employed the ridge regression best linear unbiased prediction (RR-BLUP; (Meuwissen, Hayes, and Goddard 2001; Whittaker, Thompson, and Denham 2000) model to obtain GEBVs of TBN and LA. This model equates a given trait to a linear combination of random marker effects and a random error term, as described previously (e.g., Rice and Lipka 2021), and the resulting BLUPs of each marker effect are subjected to the ridge penalty (Hoerl and Kennard 1970) during the model fitting process. The RR-BLUP model was fitted using the rrBLUP R package (Endelman 2011).

### MSTEP and USTEP models for maize

We implemented two multi-locus stepwise model selection procedures to identify markers exhibiting strong statistical associations with TBN and LA in maize. The first procedure was the multi-trait, multi-locus (MSTEP) procedure, which is described in detail in Fernandes et al. (2022). Briefly, this procedure fits a series of multi-trait, multi-locus models in a stepwise manner to identify markers exhibiting strong additive associations with multiple traits. The specific markers to be included in the model are determined through a stepwise model selection procedure. In this implementation, we considered TBN and LA as the two response variables. The second procedure we considered was a single-trait analogue of MSTEP. As done in Fernandes et al. (2022), we abbreviated this procedure as the univariate stepwise model selection procedure (USTEP), and we fitted it separately to TBN and then again to LA. For both of these model selection procedures, stepwise model selection was conducted in the TASSEL software (Bradbury et al. 2007) until a total of ten markers were in the final models.

### Genomic predictions within the Goodman-Buckler maize diversity panel

Five-fold cross-validation was performed to obtain predictive abilities for LA and TBN in the full marker set (44,930 SNPs), subsets of markers obtained from gene co-expression modules (21,362 SNPs) and from network motifs (466 SNPs) from Bertolini et al. (2024). For each of these subsets, we also evaluated the predictive abilities of markers with MAF < 0.05 and then markers with MAF > 0.05. Predictive abilities of markers selected using MSTEP and USTEP models were also included. We utilized the R packages rrBLUP (Endelman 2011) and GAPIT (Lipka et al. 2012) along with in-house developed R scripts for genomic predictions using the RR-BLUP mixed model. The predictive ability was calculated as the sample mean Pearson product-moment correlation coefficient (*r*) between observed trait values and genomic estimate breeding values (GEBVs) across all validation sets.

### Using the Goodman-Buckler diversity panel to train models for genomic prediction in the Ames maize panel

To validate the models trained using the Goodman-Buckler diversity panel on unobserved data, a subset of 1,064 individuals of the Ames panel that maximize diversity in TBN and LA (Bertolini et al. 2024) was selected for comparison of genomic predictions (GEBVs) and observed trait values for LA and TBN from this study. The various predictions were performed in the aforementioned categories of marker sets.

### Genomic prediction within the SAP in sorghum and the HDRA in rice

A five-fold cross validation procedure, very similar to that described for the Goodman-Buckler maize diversity panel, was used to evaluate the predictive ability of markers in the vicinity of sorghum and rice syntenic orthologs of the GC and NM genes. For the SAP, this resulted in a total of 59,995 markers in the vicinity of the GC orthologues and 2,695 markers in the vicinity of NM orthologues. For the HDRA, we similarly obtained a total of 293,509 markers in the vicinity of GC orthologues and 10,970 markers in the vicinity of NM orthologues.

### Procedure for obtaining an empirical null distribution to test for contribution of gene modules to genomic signals underlying traits

We followed a procedure similar to Parvathaneni *et al*. (2020) to derive an empirical distribution of prediction accuracies under the null hypothesis that the genes underlying the signals captured in the two gene set categories (i.e., GC and NM) are not important. For each category in each of the above genomic prediction experiments, we generated 1,000 random subsets by randomly selecting genes. Each of these subsets contained the same number of genes as those included in the selected category. Within each random subset, we then selected SNP markers using the genomic coordinates as described in the previous sections. We then fitted an RR-BLUP model in the respective training sets using only the SNP markers included in the random subset. Consequently, we obtained an empirical distribution of prediction accuracies under this null hypothesis. The predictive ability of the given gene set was then compared to this empirical null distribution, and a P-value was subsequently calculated. We considered statistical significance at 𝛼 = 0.05.

### Data availability

All data are publicly available. The scripts used to conduct genomic prediction will be made publicly available upon acceptance of this manuscript.

## Results

### Prediction accuracies of TBN and LA traits are enhanced by using gene network informed marker sets

To assess the ability of the NM and GC marker sets to accurately predict GEBVs of TBN and LA for 231 accessions in the Goodman-Buckler maize diversity panel (Flint-Garcia et al. 2005), we conducted a five-fold cross-validation procedure. Our results suggested that SNPs in the vicinity of GC genes had similar predictive ability to the entire whole-genome marker set (Figure 2) with a mean predictive ability of 0.60 and 0.56 for TBN and LA, respectively. The predictive abilities observed in markers around GC genes were significantly greater than those derived from markers near randomly selected genes for both traits (Figure 3). We also noted that the predictive ability of the GC set did not drop severely when using only low-MAF (MAF < 0.05) SNPs (Supplemental Table 1).

**Figure 2:**
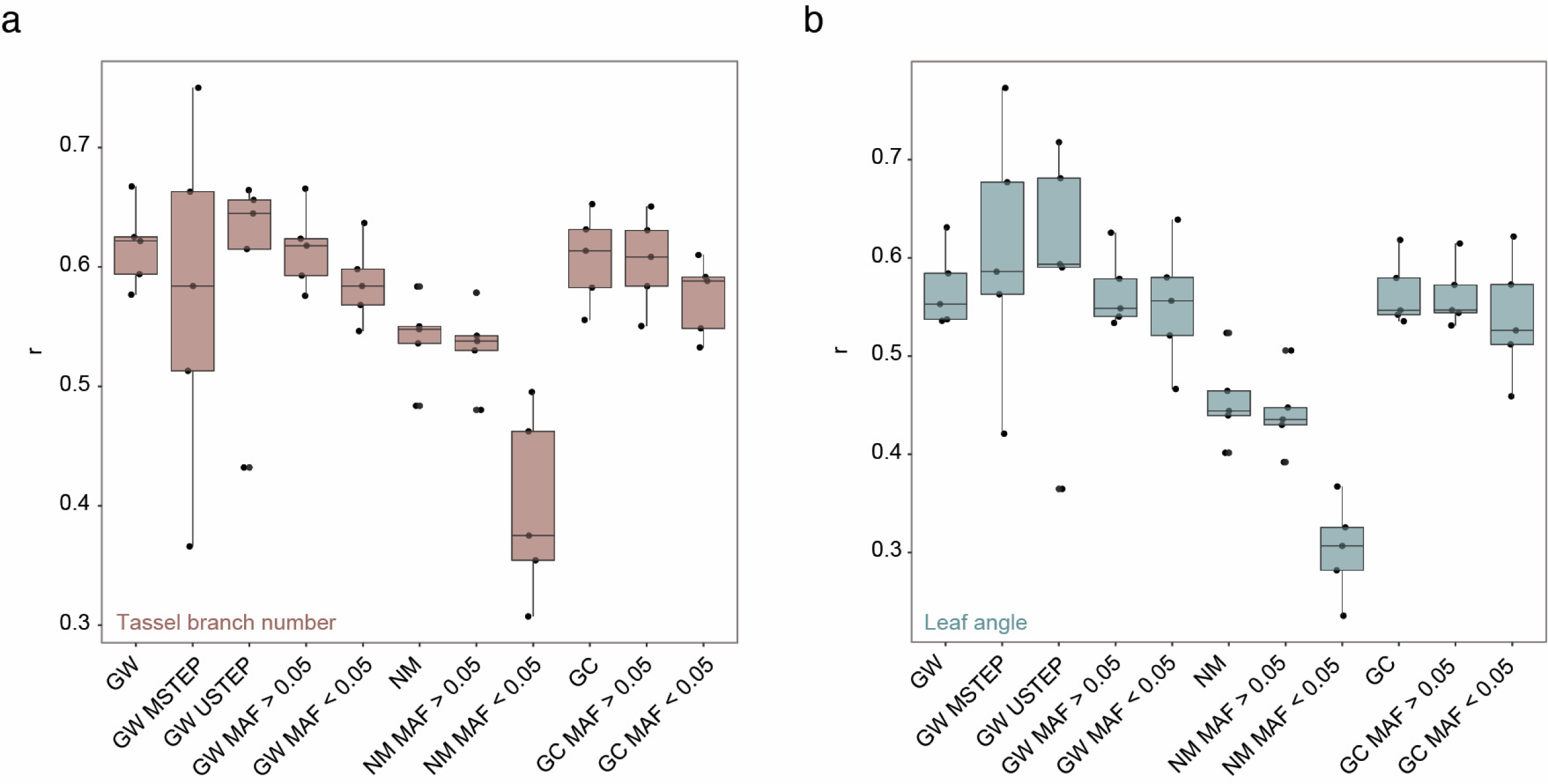
Predictive ability in the Goodman-Buckler diversity panel. The image depicts the prediction accuracies in the training set. The boxplots illustrate the results of the five-fold cross-validation tests conducted on the genome-wide (GW), the network motif (NM), and the gene co-expression (GC) marker sets at different minor allele frequencies (MAF) cut-off. Y-axis represents correlation coefficient (r) between observed trait values and GEBVs; panels **a** and **b** show results on TBN and LA, respectively.

**Figure 3:**
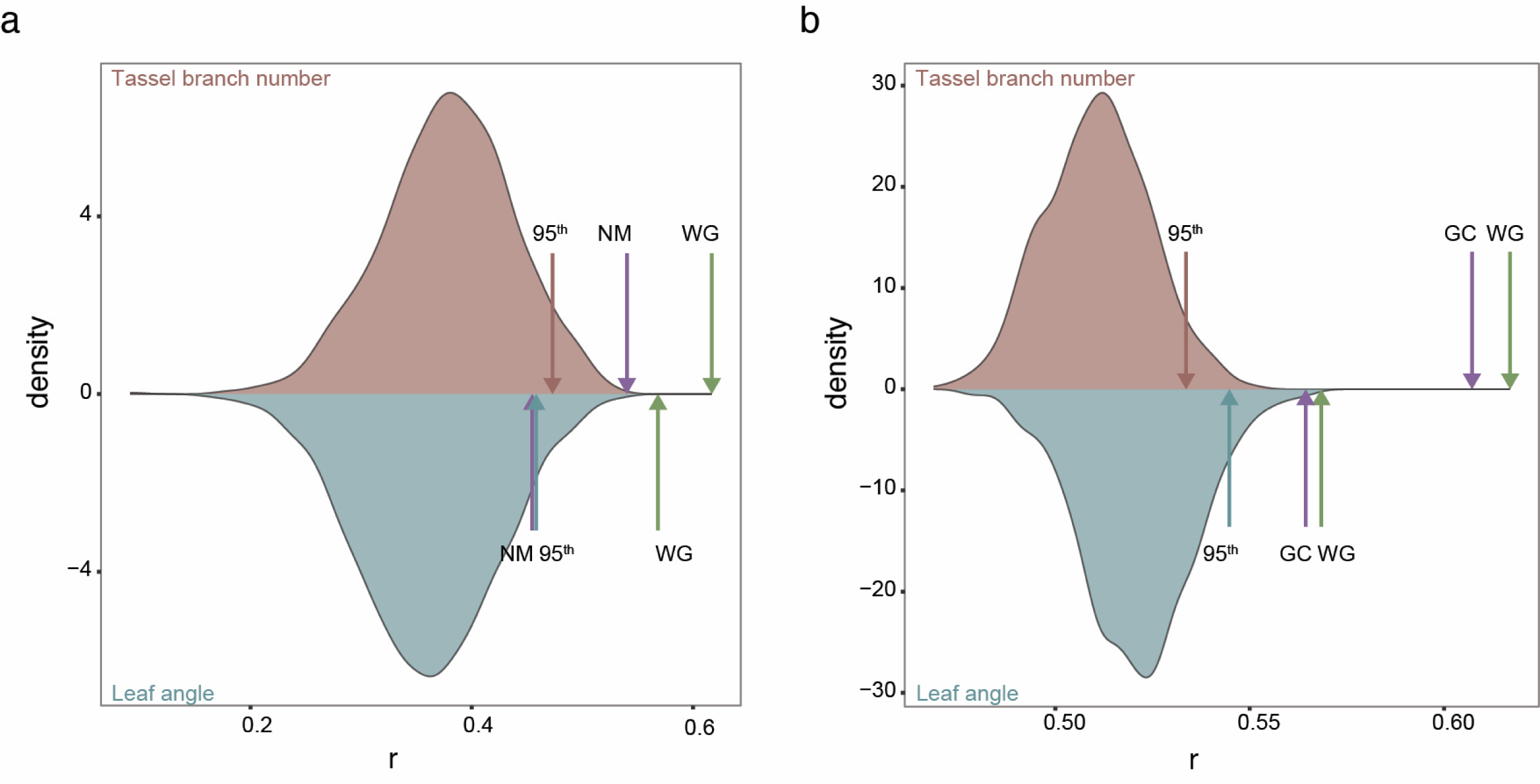
Predictive performance of network motif and gene co-expression marker sets in the training set. The density plots illustrate the empirical null distributions, above and below zero density, generated based on 1,000 iterations of randomly selected genes and the subsequent selection of the co-localizing markers. Prediction results for TBN and LA are represented in mauve and teal, respectively, for the network motif (NM) set (panel **a**) and the gene co-expression (GC) set (panel **b**). Arrows indicate the 95^th^ percentile of the empirical null distributions, the five-fold cross predictive ability (average) for the NM and GC set (purple) and the whole genome set of markers (green) conducted using the Goodman-Buckler diversity panel.

Notably, markers associated with NM genes, despite accounting for only 10% of the whole-genome SNPs, showed only slightly lower predictive abilities for both traits (Figure 2) relative to the whole-genome markers. When compared to the respective empirical null distribution, we observed that markers near the NM genes showed greater ability to predict TBN than was expected under the null hypothesis. In contrast, the predictability of LA by markers near the NM genes was not substantially greater than the range of prediction accuracies observed across the corresponding empirical null distribution of prediction accuracies (Figure 3). For both traits, the predictive ability of the markers identified by MSTEP and USTEP were similar to those from the genome-wide marker set. However, the results suggested that relative to a genome-wide set of markers, MSTEP and USTEP have potential to increase the variability (and hence uncertainty) in prediction accuracies.

### Markers near NM genes capture unique genomic signals in the Ames diversity panel

We next assessed the ability of GP models trained in the Goodman-Buckler diversity panel to predict GEBVs in the Ames panel. The predictive ability of GC markers was 0.55 for TBN and 0.46 for LA, while the whole-genome set achieved an accuracy of 0.57 and 0.47, respectively, for TBN and LA (Figure 4). Interestingly, we noted that low-MAF GC markers yielded identical LA predictive abilities as the low-MAF subset of whole-genome markers, but not for TBN. We observed that the predictive abilities of markers near GC genes exceeded those of markers near NM genes in the Ames panel (Supplemental Table 1 and Figure 4). These outcomes closely mirrored those observed during cross-validation experiments conducted within the Goodman-Buckler diversity panel (Supplemental Table 1). However, when compared to their respective empirical null distribution of predictive abilities of markers near randomly selected genes, the predictive ability of the markers near GC fell below the 95^th^ percentile (Figure 5a). In comparison, the predictive ability of NM genes were substantially higher than what would be expected under the null hypothesis that the NM genes are not making a meaningful contribution to the genomic signal of TBN and LA (Figure 5b). Finally, we observed that the predictive abilities of the markers identified from MSTEP and USTEP in the Goodman-Buckler diversity panel were notably lower than those for any other considered subset of markers.

**Figure 4:**
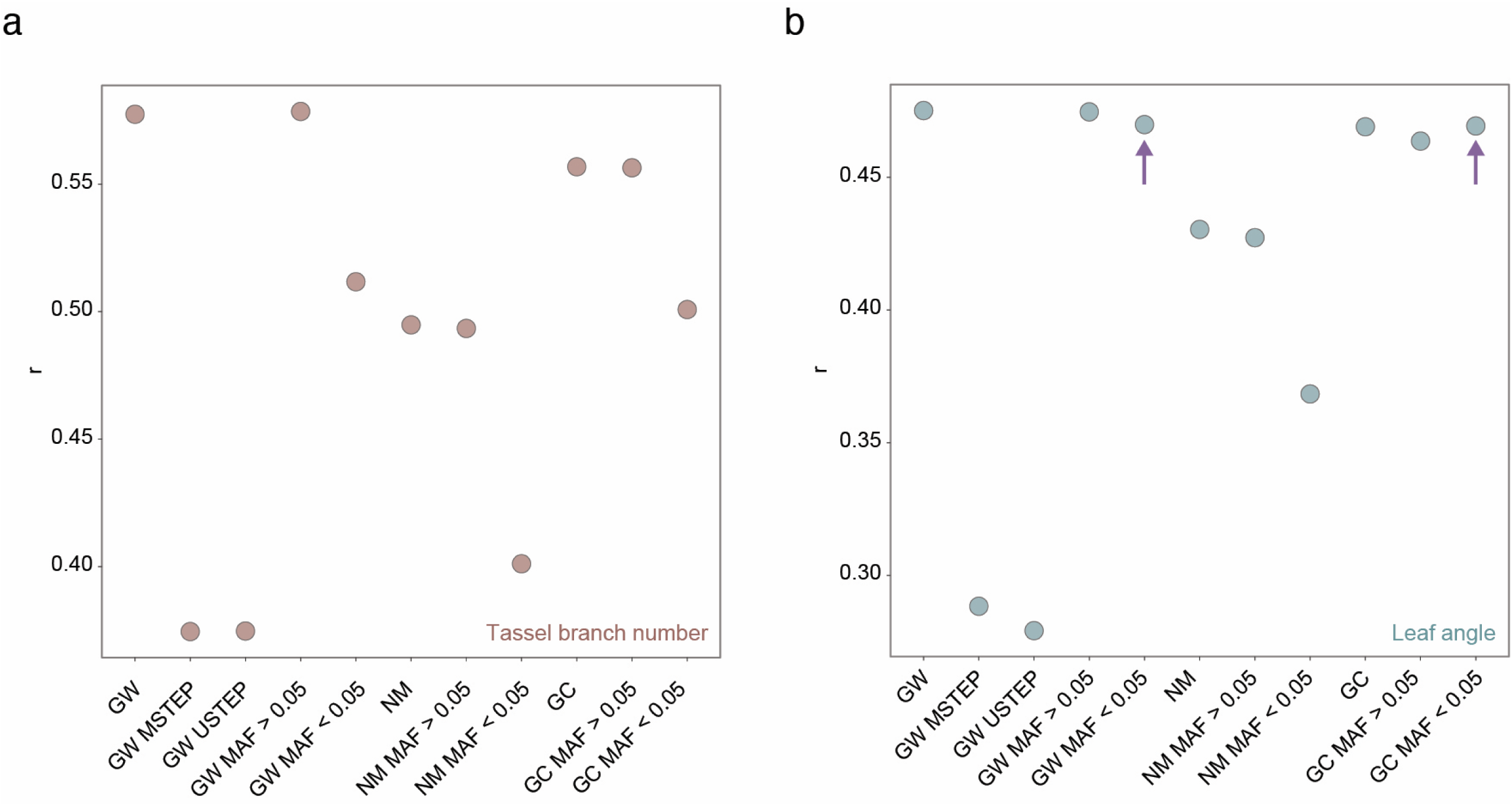
Predictive ability of network motif and gene co-expression marker sets in the validation set. The image represents the predictive ability results in the validation set for TBN (**a**) and LA (**b**) in the Ames diversity panel. Each dot represents the correlation coefficient (r) of different maker sets: genome-wide (GW), network motif (NM) and the gene co-expression (GC) markers at different minor allele frequencies (MAF) cut-off.

**Figure 5:**
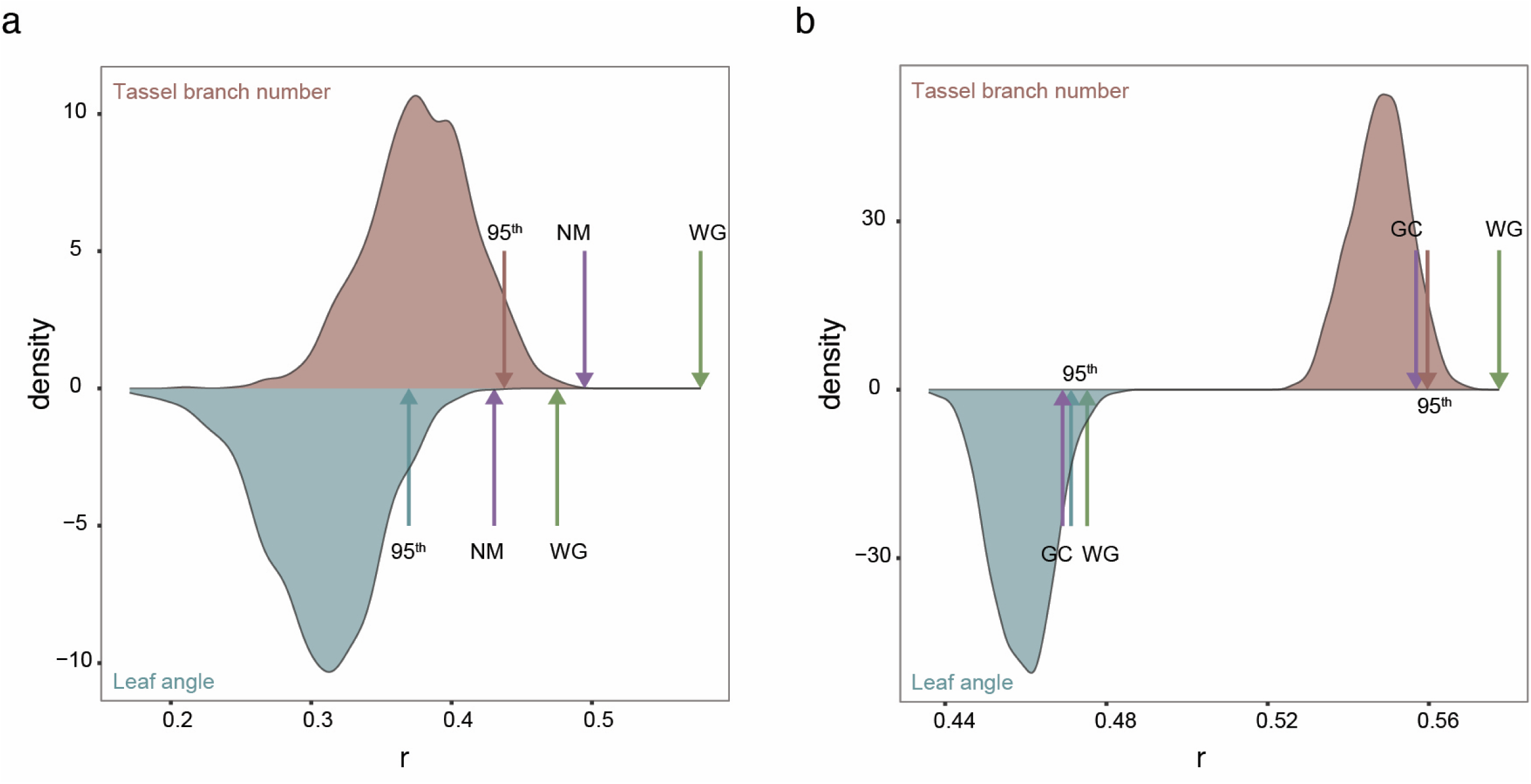
Predictive performance of network motif and gene co-expression markers in the validation set. The density plots illustrate the empirical null distributions, above and below zero density, generated based on 1,000 iterations of randomly selected genes and the subsequent selection of the co-localizing markers. Prediction results for TBN and LA are represented in mauve and teal, respectively, for the network motif (NM) set (panel **a**) and the gene co-expression (GC) set (panel **b**). Arrows indicate the 95^th^ percentile of the empirical null distributions, the predictive ability for the NM and GC set (purple) and the whole genome set of markers (green) conducted using the Ames diversity panel. Δ represents the difference in predictive ability between the WG and the marker subsets, NM or GC.

### Markers near NM orthologs in sorghum showed higher predictive abilities for LA than expected by chance

To test whether context-specific biological data from maize could be used for accurately predicting parallel phenotypes in sorghum, a closely related cereal crop, we used the GC and NM genes in maize to predict LA in sorghum. We observed that the ability of markers proximal to sorghum syntenic orthologs of GC genes to predict GEBVs of LA was similar to those in the vicinity of orthologous NM genes (Supplemental Table 2 and Figure 6), with a mean predictive ability of 0.50 and 0.51 for GC and NM, respectively. However, we also noted that the empirical null distribution of predictive abilities from randomly selected genes corresponding to the GC set tended to be larger than a comparable distribution corresponding to the NM set (Figure 6). These results suggest that markers associated with NM genes play a more substantial role in the genomic signal underlying LA variance in the SAP. These overall findings closely match those from the analysis of the maize lines in the Ames diversity panel.

**Figure 6:**
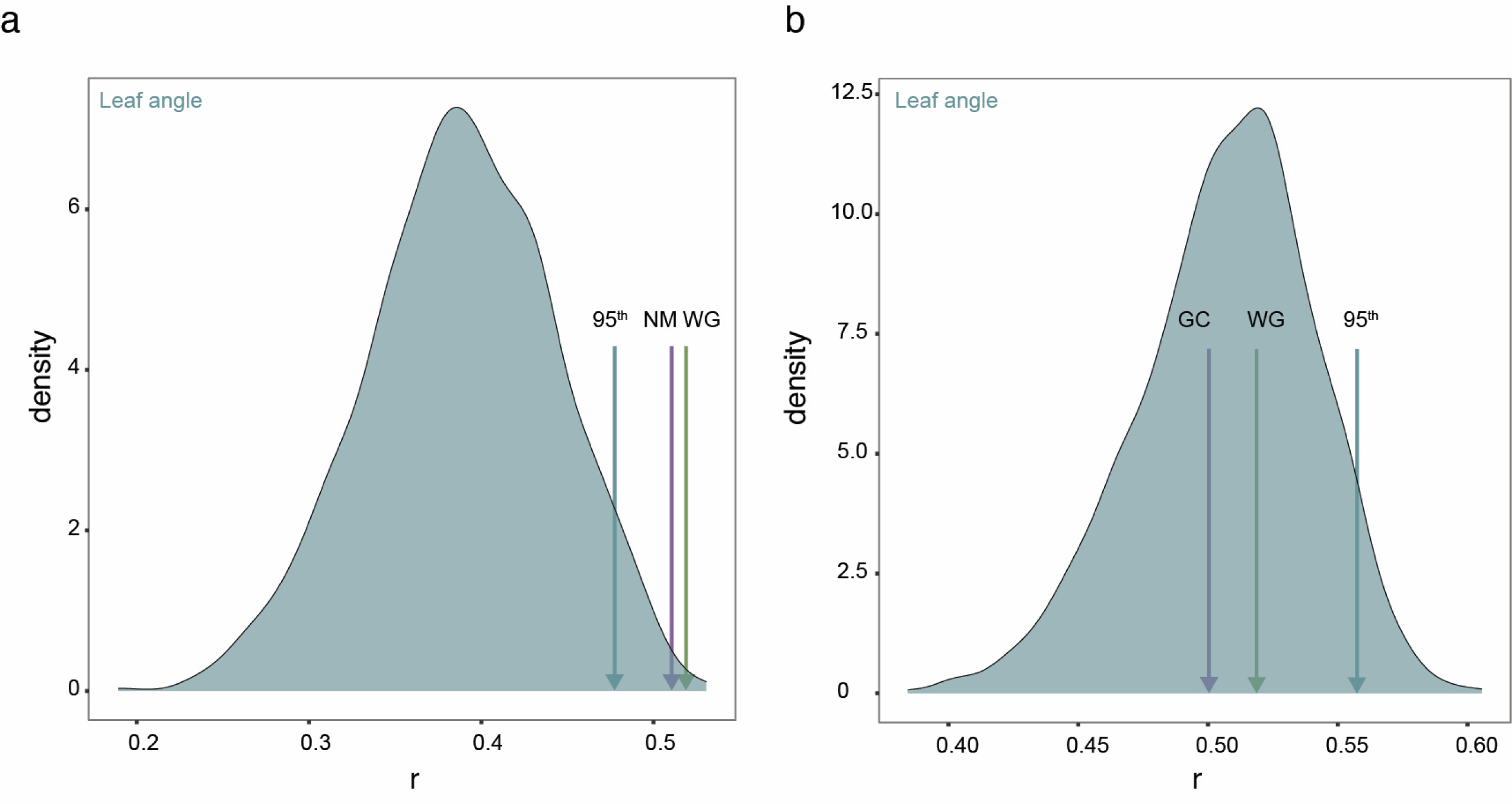
Cross-species predictive ability between maize and sorghum. The density plots illustrate the empirical null distributions generated based on 1,000 iterations of randomly selected sorghum genes and the subsequent selection of the co-localizing markers. LA prediction results are represented in teal both for the network motif (NM) set (panel **a**) and the gene co-expression (GC) set (panel **b**). Arrows indicate the 95^th^ percentile of the empirical null distributions, the five-fold cross predictive ability (average) for the NM and GC set (purple) and the whole genome set of markers (green) conducted using the Sorghum Association Panel.

### Cross-species predictive ability for LA between maize and rice is not statistically significant

We further assessed the cross-species translatability of markers associated with network genes and their applicability between more distantly related grass species, i.e., maize and rice. As done in sorghum, we predicted GEBVs of rice LA based on SNPs near rice syntenic orthologs of GC and NM genes. The whole-genome marker set yielded a mean predictive ability of 0.27. Given the low trait heritability (*h^2^*= 0.32, see Huber et al., 2023) this level of prediction was expected. However, the predictive abilities of the GC and NM sets tended to be less than those from selected markers used to generate the null distribution (Supplemental Table 3 and Figure 7).

**Figure 7:**
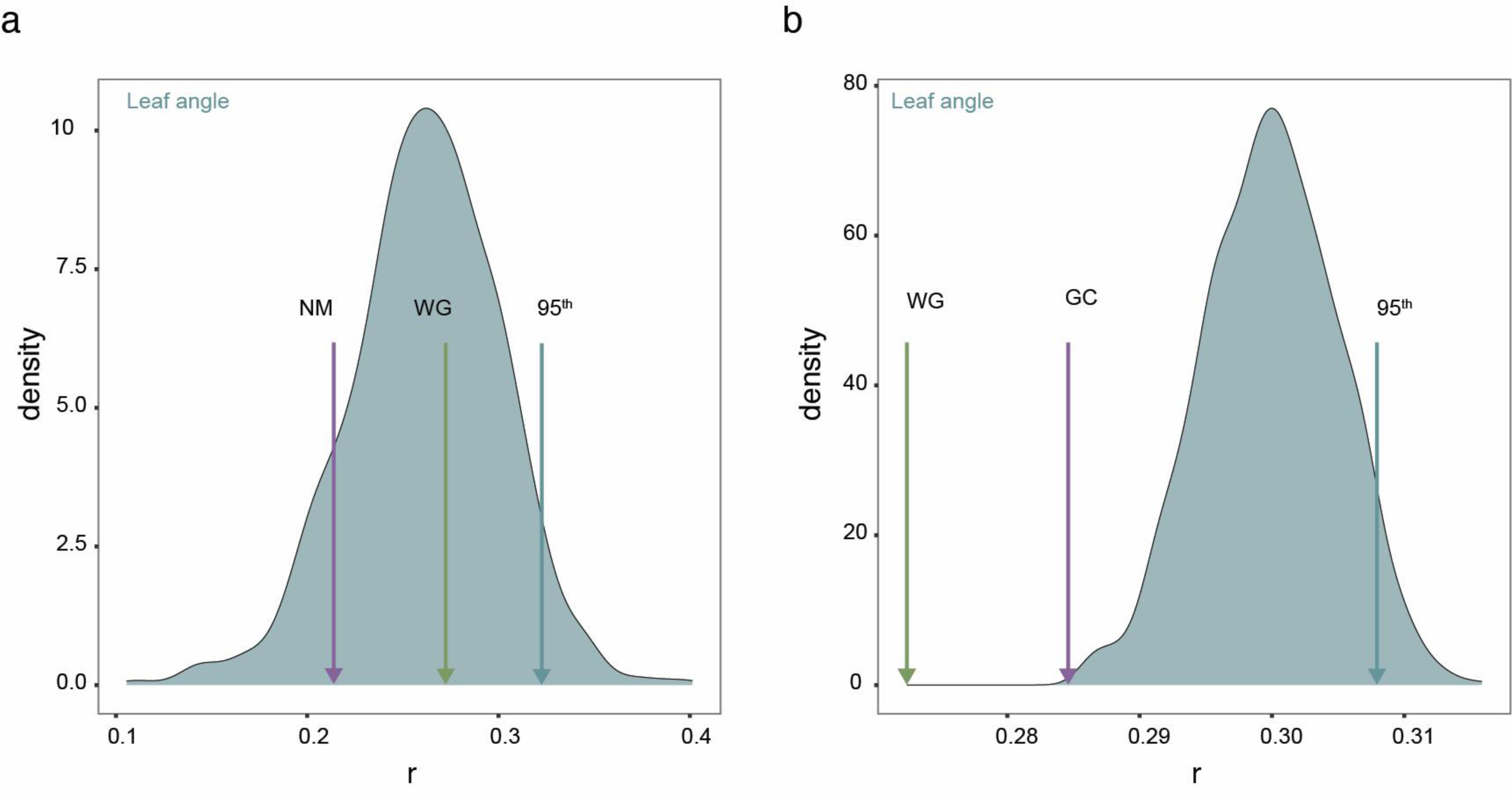
Cross-species predictive ability between maize and rice. The density plots illustrate the empirical null distributions generated based on 1,000 iterations of randomly selected rice genes and the subsequent selection of the co-localizing markers. LA prediction results are represented in teal both for the network motif (NM) set (panel **a**) and the gene co-expression (GC) set (panel **b**). Arrows indicate the 95^th^ percentile of the empirical null distributions, the five-fold cross predictive ability (average) for the NM and GC set (purple) and the whole genome set of markers (green) conducted using the High-Density Rice Array panel.

## Discussion

We assessed the ability of markers located within and around two biological network-informed gene sets from maize to predict breeding values of plant architecture traits in three agronomically important crops. We observed that the set of transcription factor-encoding genes associated with recurrent NMs gave higher predictive abilities in maize and sorghum than expected by chance, but not in rice. This suggests that regulatory networks derived from one species (i.e., maize) can be used to inform loci contributing to the genetic architecture in a closely related species (sorghum). Our results also showed that this did not hold up when translating to rice, a more distantly related species; however there are other factors that may have confounded this analysis, as described below.

### Network motifs can capture information underlying the genetic architecture of LA important for cross-species inferences

Our results support our hypothesis on the contributions of context-specific gene regulatory networks to the genetic architecture of LA and TBN. This suggests that finely-tuned GP models including only markers in the vicinity of NM genes can effectively infer elements of the genetic architecture of complex traits. Results from our cross-species analyses are likely attributed to a shared set of functionally constrained regulators that play important roles in the genotype to phenotype relationship underlying LA genetic architecture in maize and sorghum, but not in rice. This could be indicative of the close evolutionary distance between maize and sorghum (Wang et al. 2015) and aligns with prior findings showing gene regulatory conservation among syntenic orthologs in these species (Zhang et al. 2017).

However, in the case of rice we must account for differences in the developmental stage at time of phenotype collection (these were collected at an early vegetative growth stage), enhanced tillering compared to the other species, and the different method used for quantifying LA in the rice data set (Huber *et al*. 2023). All of these factors may have confounded our results. Ideally, GP models in rice should be trained on LA data from mature rice plants as was done for maize and sorghum. Accounting for these differences could rule out the possibility that the observed low prediction accuracies for markers near synthenic rice orthologs of maize NM features arose because the key transcription factors underlying LA in rice change across growth stages.

### Gene co-expression networks sufficiently capture meaningful contributions to genetic architecture within the panel where the traits were quantified

In contrast to our results with the NM gene sets, the only situation where we saw evidence of markers surrounding GC network genes yielding higher prediction accuracies than expected by chance was within the Goodman-Buckler diversity panel. This suggests that the causal variability captured by the GC networks could be very specific to the data set being analyzed, and is potentially prone to overfitting to such an extent that they cannot capture signatures of genetic architecture, even for different panels within the same species. More broadly, this result implies that using correlations between gene expression is insufficient for capturing markers that have high accuracy when predicting unrelated individuals. This could reflect the co-expression network rewiring influenced by specific selective pressures. Network motifs instead capture building block patterns within the complex networks of recurrent transcription factors that might preserve functional conservation intra/inter species. Therefore, the identification of recurring transcription factors associated with three-node network motifs helps prioritize genes that are more likely to serve as key regulators, as well as subset markers linked to genes that contribute to the genetic architecture in unrelated individuals and environments.

### Using predictive abilities from GP to infer genetic architecture

This study demonstrated the potential for using GP to make inferences on genetic architecture. In practice, GP is almost overwhelmingly used to predict GEBVs of crops or livestock using whole-genome marker data, as opposed to inferring which genomic regions are likely to contain features that biologically control trait variability (Rice and Lipka 2021).Indeed, the application of GP to make such inferences should be discouraged because the number of markers in a typical data set vastly exceeds the number of individuals (see de Los Campos et al. 2013) for an overview of genomic prediction models). This “p >> n” scenario leads to the priors and/or penalties used in GP having such a major influence on marker effect estimates that different penalties and/or priors could identify different regions of strong statistical associations for the same trait (see e.g., Gianola 2013) for an in-depth description). Similar to other studies (e.g., Turner-Hissong et al. 2020), we circumvented this problem by comparing the predictive abilities of several GP models, each that focus on biologically-informed subsets of the genome. If follow-up studies confirm our findings on the predictive abilities of the NM sets, they would underscore that substantial insight into genomic architecture can be made fitting off-the-shelf GP models to *a priori* biologically-informed marker subsets.

## Conclusion

We used an innovative GP approach informed by gene regulatory circuitries to study the genetic architecture of complex traits. Our analyses suggest that NM facilitates the translation of biological information related to plant architecture across closely related species, such as maize and sorghum. This suggestive convergence of functionally constrained regulators underlying the genetic architecture of LA opens up promising avenues for targeted breeding practices in sorghum, which can lead to optimized plant architecture for high-density planting and enhanced agricultural productivity.

## Acknowledgements

We acknowledge the National Center of Supercomputing Applications at UIUC for the computational facilities that made it possible to conduct the analyses in a timely manner.

